# *In-vivo* antidiarrheal activity: From the crude extract and solvent fractions of *Rhamnus prinoides* (Rhamnaceae) leaves

**DOI:** 10.1101/2022.09.19.508518

**Authors:** Teklie Mengie Ayele, Endeshaw Chekol Abebe, Zelalem Tilahun Muche, Melaku Mekonnen Agidew, Yohannes Shumet Yimer, Getu Tesfaw Addis, Nega Dagnaw Baye, Achenef Bogale Kassie, Muluken Adela Alemu, Tesfagegn Gobezie Yiblet, Gebrehiwot Ayalew Tiruneh, Samuel Berihun Dagnew, Tilaye Arega Moges, Tesfaye Yimer Tadesse, Amein Ewnetei Zelalem

## Abstract

**Objectives:** The inherent toxicities of the drugs urge the search for alternative drugs that are safe and effective. Therefore, the objective of the study is to evaluate the *in-vivo* anti-diarrheal activity of crude extract and solvent fractions of *Rhamnus prinoides* leaves.

**Methods:** The Leaves of *Rhamnus prinoides* were macerated using absolute methanol and then fractionated. For *in-vivo* antidiarrheal activity evaluation of the crude extract and solvent fraction, castor oil-induced diarrhea, castor oil-induced anti-enteropolling, and intestinal transit models were used. One-way analysis of variance was used to analyze the data, followed by a Tukey post-test. The standard and negative control groups were treated with loperamide and 2% tween 80 respectively.

**Results:** A significant reduction in the frequency of wet stools and watery content of diarrhea, intestinal motility, intestinal fluid accumulation, and delaying the onset of diarrhea as compared with controls were observed in mice treated with 200 mg/kg and 400 mg/kg ME. However the effect increased dose-dependently, and the 400 mg/kg ME produced a comparable effect with the standard drug in all models. Amongst the solvent fractions, n-BF significantly delayed the time of diarrheal onset and reduced the frequency of defecation, and intestinal motility at doses of 200mg/kg and 400mg/kg. Furthermore, the maximum percentage inhibition of intestinal fluid accumulation was observed in mice treated with 400 mg/kg n-BF (p<0.01; 61.05%).

**Conclusion:** The results of this study showed that crude leaves extract and solvent fractions of *Rhamnus prinoides* had significant anti-diarrheal activity, providing scientific support for its traditional use as a diarrhea treatment.

## Introduction

More than 80% of the world’s population, according to the World Health Organization (WHO), relies on traditional medicine for their primary healthcare requirements. It is commonly stated that plants are responsible for 25% of all medications given today. Plant-derived medications, according to this estimate, account for a large portion of natural product–based pharmaceuticals [1]. Herbal products derived from medicinal plants are chosen because they require less testing, are safer, more efficient, culturally acceptable, and have fewer adverse effects [2].

There have been numerous reports of traditional plants being used to treat diarrheal illnesses. Herbal goods are believed to be more compatible with the human body since they contain chemical substances that are a part of organism physiology [2]. Plant extracts can lower electrolyte release, delay gastrointestinal transit, restrict gut motility, enhance water adsorption, and lessen spasms [3]. These activities may help to explain why certain plants are beneficial in the treatment of diarrheal disease.

Although a large portion of the world’s population relies on traditional medicine for primary health care, traditional medications and medical techniques are passed down verbally through generations, and in most cases, the effective doses and combinations proposed by traditional healers differ; as a result, the effective doses, as well as the effectiveness, safety, toxicity, and chemical composition variation between plant parts, are not fully understood [4].

Furthermore, antimicrobial-resistant strains have become widespread, posing a global danger to currently available antimicrobial ineffective medicines, particularly antidiarrheal drugs. According to the WHO’s 2014 global report on antimicrobial resistance, bacteria that cause diarrhea, such as *E.coli* (to 3rd generation cephalosporins, and fluoroquinolones), *Neisseria gonorrhoeae* (to 3rd generation cephalosporins), and Shigella species, have decreased susceptibility to fluoroquinolones. Apart from this, the majority of currently available medications have side effects such as bronchospasm, vomiting (racecadotril), intestinal blockage, constipation (Loperamide), and addiction [5]. As a result, it is a must to expand research on culturally preserved medicinal plants to discover alternative pharmaceuticals derived from natural sources.

Furthermore, there is a wealth of ethnomedical information on plant usage in the scientific literature that has yet to be gathered in a useable manner [1]. *Rhamnus* prinoides (*R. prinoides*) (Rhamnaceae) is also known as Gesho in Ethiopia. It is used in traditional medicine to treat headaches, neck ulcers, and edema [6]. It is one of the numerous herbs that have traditionally been used to treat diarrhea. This study will confirm the plant’s traditional use and elucidate the nature of the phytochemical constituents responsible for its effect and possible ways of antidiarrheal action. The results from the current study can be used to aid in the development of novel anti-diarrheal agents that address issues with current anti-diarrheal medications. It will also guide traditional users on how to prepare and use the plant in various ways. Furthermore, the findings of this study will aid the scientific community in furthering their research into the plant *R. prinoides* by launching advanced studies into molecular mechanisms and formulation of plant source drugs, as well as identifying the specific agent responsible for the anti-diarrheal effect.

Phytochemical screening revealed the presence of phenols, flavonoids, alkaloids, saponin, glycoside, and tannins in this medicinal plant, as well as the presence of five different labdane types of diterpene lactones in Ethiopian collections of *R. prinoides* leaves [7]. Furthermore, *R. prinoides* have been shown to have analgesic, anti-inflammatory, antihelmintic, and antibacterial action in animals [6]. In multiple experimental models, these findings suggest that *R. prinoides* may have anti-diarrheal potential. As a result, the aim this study is to prove the reputed antidiarrheal action of the leaves crude extract and solvent fractions *R. prinoides* based on ethno medicinal use in Ethiopia.

## Materials and Methods

### Drugs and chemicals

In this study, we used castor oil from Amman Pharmaceutical Industries in Jordan, charcoal from Acura Organics Ltd in New Delhi, tween 80 from Uni-Chem in India, loperamide hydrochloride from Medichemie Ltd in Cyprus (EU), misoprostol from Mylan Laboratories in India, distilled water from Debre Tabor University’s Department of Chemistry, n-butanol from Chloroform, methanol 99.8% from Blulux, India. The reagents were all of an analytical caliber.

### Plant materials

The leaves of *Rhamnus prinoides* (Rhamnaceae) **(Figure 1)** were collected from Debre Tabor Town, South Gondar, Northcentral Ethiopia. After collecting the plants, identification and authentication of the plant’s specimens were done by taxonomists at the Department of Biology, College of Natural Sciences and Computation, Debre Tabor University, and the voucher number TMA001/2022 with specimens were deposited for future reference.

**Figure 1:**
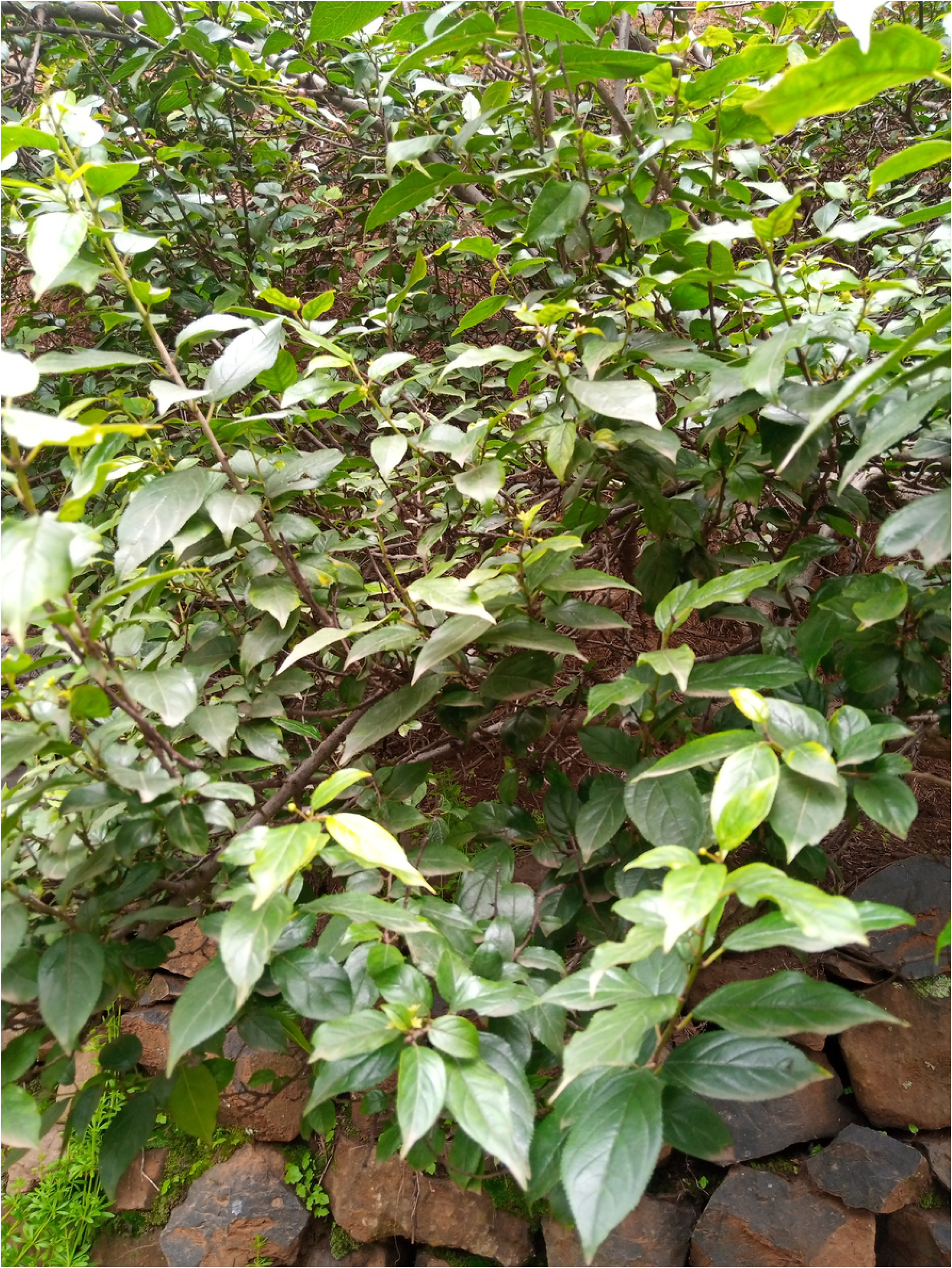
Picture of *R. prinoides* from the site of collection.

### Experimental animals

Mice of both sexes (20-30g) were obtained from the Ethiopian Public Health Institute’s animal house section in Addis Ababa. The mice were kept in polypropylene cages under conventional environmental conditions, with a 12-hour light-dark cycle and unlimited pellet food and drink. Before starting the experiment, the animals were given a week to acclimate. During the light period, all experiments were carried out. The Ethical clearance and permission were accepted and obtained from the Debre Tabor University Research and Ethical Review Committee. All the protocols were carried out following the guideline for the care and use of laboratory animals [8].

### Preparation of crude extract and solvent fractions

To eliminate dust elements, the dried leaves of *R. prinoides* were first washed in distilled water. Before extraction, it was ground with a grinder and roughly powdered with a mortar and pestle. Seven hundred grams of powdered leaves of *R. prinoides* were macerated using one liter of absolute methanol (1:6 w/v). The mixture was then filtered through Whatman No. 1 filter paper, and the residue was re-macerated with fresh solvent exhaustively. The filtrates from the three batches were mixed and concentrated in a rotary evaporator at 40°C to remove methanol. After drying, black sticky residue weighing 90 grams (12.86%) of *R. prinoides* crude extract was collected.

In a separatory funnel, 180 ml of n-butanol was used to suspend 80 grams of the crude extract, and then an equal amount of petroleum ether was added and thoroughly combined. After allowing the mixture to separate into a discrete layer, the petroleum ether portion was isolated by eluting the bottom layer. The leftover material was then combined with an equivalent amount of chloroform and separated in the same way. The solvent fractions were all concentrated and dried in a rotary evaporator at 40 degrees Celsius after that. From the dried fractions, n-butanol, chloroform, and petroleum ether had respective yields of 32.5 g (40.63%), 29 g (36.25%), and 18.5 g (23.19%). Finally, the dried fractions were reconstituted with 2% tween 80 and kept at −20°C till commencing the experiment.

### Acute Toxicity Test

The test was initially carried out utilizing the limit test criteria of the Organization for Economic Cooperation and Development (OECD) 425 Guideline (OECD, 2008) [9]. A single female mouse was administered the tested dose of 2000 mg/kg crude extract and solvent fractions as a single dose through oral gavage for a sighting study in order to establish the starting dose. Four additional female mice were used and given the same amount of the test chemicals because there was no sign of death after 24 hours. The mice were monitored for general signs and symptoms of physical and behavioral toxicity every 4 hours at a 30-minute interval for 14 consecutive days at a 24-hour interval. Then, one-tenth of 2000 mg/kg (200 mg), half of one-tenth (100 mg), and twice one-tenth (400 mg) dosages were employed for the main study.

### Grouping and Dosing

For the ME and solvent fractions, mice of either sex (weighing 20–30g) were randomly allocated into five and eleven groups, each with six animals. Prior to the test, the mice fasted for 18 hours while having unlimited access to water. The first two groups were given 10 ml/kg of 2 percent tween80 as a negative control and 3 mg/kg of loperamide as a positive control, respectively, for the investigation of the antidiarrheal properties of both ME and the solvent fractions. The ME and two solvent fractions of R. prinoides leaves were administered to the remaining groups at test doses of 100, 200, and 400 mg/kg. In all current models, diarrhea was induced using castor oil. All of the doses were administered orally.

### Phytochemical screening of the crude extract and solvent fractions

Standard tests were used to conduct preliminary phytoconstituent screening of secondary metabolites from both ME and the solvent fractions of the leaves of *R. prinoides* [10, 11].

#### Test for saponins

The 5 ml distilled water was added to 0.25 grams of ME and the solvent fractions of the leaves of *R. prinoides*. The solution was then violently shaken, and stable, continuous foam was seen. Saponins were detected by the formation of a steady froth that lasted about half an hour.

#### Test for terpenoids

The 2 ml chloroform was added to 0.25 grams of ME and the solvent fractions of the leaves of *R. prinoides*. Then, to build a coating, 3ml concentrated sulfuric acid was carefully applied. The presence of terpenoids was indicated by a reddish brown coloration of the interface.

#### Test for tannins

In a test tube, 0.25 grams of ME and the solvent fractions of the leaves of *R. prinoides* were cooked in 10 ml water and then filtered using filter paper (Whatman No. 1). To the filtrate, a few drops of 0.1 percent ferric chloride were added. Brownish green or blue-black precipitate is a sign of tannin content.

#### Test for flavonoids

10 ml of ethyl acetate, 0.2 grams of ME, and the solvent fractions of R. prinoides leaves were heated in a water bath for 3 minutes. The mixture was cooled after filtering. The filtrate was then combined with 1 ml of a mild ammonia solution in 4 ml and agitated. When the layers were allowed to separate, the ammonia layer’s yellow tint confirmed the presence of flavonoids.

#### Test for cardiac glycosides

The 2 ml glacial acetic acid containing one drop of ferric chloride solution was added to 0.25 grams of ME and the solvent fractions of the leaves of *R. prinoides* that had been diluted with 5 ml of water. The 1 ml of sulfuric acid was used as a base. The presence of a deoxysugar, which is characteristic of cardenolides, was identified by a brown ring at the interface.

#### Test for steroids

The 0.25 grams of ME and the fractions of the leaves of *R. prinoides* were mixed with 2 ml sulfuric acid and two ml acetic anhydride. Some samples changed color from violet to blue or green, indicating the presence of steroids.

#### Test for alkaloids

A few drops of freshly made Mayer’s reagent were added to 0.5 grams of ME and the solvent fractions of the leaves of *R. prinoides*. The presence of alkaloids was determined by the production of the cream.

### Determination of anti-diarrheal activity

#### Castor oil-induced diarrhea model

To test the anti-diarrheal properties of the ME and the solvent fractions of the leaves of *R. prinoides*, we used the castor oil-induced diarrheal model explained by Umer *et al*. (2013) with minimal modification [12]. Mice of either sex fasted overnight were divided into five and eleven groups (6 mice per group) for the crude extract and solvent fractions respectively and (90). As detailed in the grouping and dosing section, the mice received the given doses. Each mouse received castor oil, 0.5 ml orally 1hour after treatment. The mice were then kept in separate cages with the bottoms wrinkled with white paper so that the number and consistency of feces could be examined. The papers were changed every l hour to allow the feces to be seen and counted, as well as to ensure stool consistency. Normal pelleted feces (0), distinct soft-formed feces (1), soft-formed feces (2), soft watery stool (3), and watery stool with minimal solid substance (4) were used to evaluate diarrhea (4) [13].

The mice were monitored for 4 hours, during which the amount of dry and wet feces excreted by the mice was tallied and compared to the negative and positive controls to determine the antidiarrheal activity of the ME and the solvent fractions of the leaves of *R. prinoides*.

The time period designated as the onset is the time interval in minutes between the application of castor oil and the appearance of the first feces. For the negative control, the total amount of feces was assumed to be 100 percent, and the percentage of diarrheal inhibition for moist and watery content of feces was calculated using the following formula:

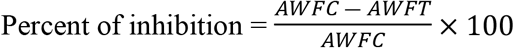

‘AWFC’ stands for the average weight of the fecal matter of controls and ‘AWFT’ stands for the average weight of fecal matter of test groups [13].

### Castor oil-induced enteropooling

The method previously mentioned by Amuzat *et al*. (2020) was used to investigate the effects of ME and the solvent fractions of the leaves of *R. prinoides* on the intraluminal fluid buildup [14]. Before the experiment, mice of either sex were fasted overnight and then randomly divided into five and eleven groups (6 mice per group) for ME and solvent fractions respectively. Then, the mice were treated with 10 ml/kg 2% tween80, 100, 200, and 400 mg/kg of ME and solvent fractions, and a standard medication (loperamide hydrochloride 3 mg/kg) orally. After 1 hour of treatment administration, the animals were given castor oil, 0.5 ml. The mice were cervically dislocated one hour after receiving castor oil.

The small intestine was then taken from each animal’s abdomen and knotted with thread at the pyloric end and the ileocaecal junction. The contents of the dissected small intestine were milked into a graduated tube and their volume was determined after the dissected small intestine was weighed. After milking, the intestine’s weight was determined, and the difference between the weight of the intestines when they were full and empty was computed. Finally, using the calculations below, the percentage inhibitions of intestinal secretion (volume and weight) were estimated relative to the negative control.

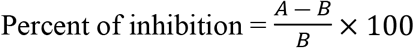

‘A’ represents the average volume or weight of the intestine in the control group, while ‘B’ represents the average volume or weight of the intestine in the test groups [12].

The weight was recorded as (m1-m0) g and the volume of the intestinal contents was read from the graduated measuring cylinder.

### Castor oil-induced gastrointestinal motility test

The method described by Uzuegbu *et al*. (2021) [15], was used with little modification to investigate the effect of the ME and the solvent fractions of the leaves of *R. prinoides* on the gastrointestinal motility. Mice of either sex fasted overnight were divided into five and eleven groups (6 mice per group) at random for crude extract and solvent fractions respectively [13]. Each animal received 0.5 ml of castor oil orally 1 hour after dosing the test agents. All mice were given 1ml of 10% charcoal suspension orally after 1 hour of castor oil administration and slaughtered after 30 minutes. The length of the small intestine was measured after it was dissected, and the distance traveled by charcoal from the pylorus to the caecum was estimated. Each mouse’s intestine was preserved in formalin to stop peristalsis, and then rinsed in distilled water before the distance covered by the charcoal was measured. The peristaltic index (PI) for the charcoal movement was calculated as follows:

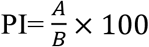

Where ‘A’ is the length of the entire intestine and ‘B’ is the distance covered by charcoal. The following formula was used to calculate the percentage of inhibition:

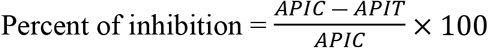

‘APIC’ stands for average PI of the control group, while ‘APIT’ stands for average PI of the test group [13].

### In vivo anti-diarrheal index (ADI)

Data from all antidiarrheal models used in this investigation were utilized to compute the ADI of treatment groups using the formula below [14].

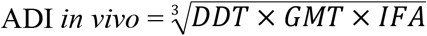

Where ‘DDT’ stands for defecation delay time (as a percent of control) and the gastrointestinal motility (GMT) is affected by a reduction in charcoal transport (as a percent of control, IFA is the decrease in the intestinal fluid accumulation (as % of control).

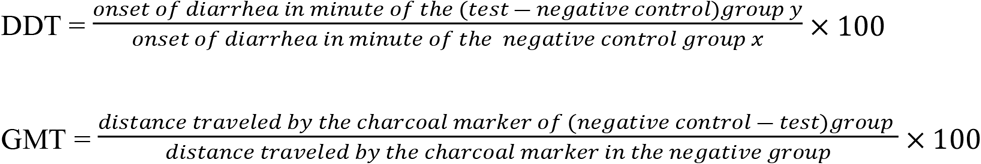

### Statistical Analysis

The mean and standard error of the mean is used to express the findings (SEM). The current study’s experimental data were evaluated with the software Statistical Package for Social Sciences (SPSS), version 20, and statistical significance was assessed using one-way analysis of variance (ANOVA) and the Tukey Kramer posthoc test. A statistically significant P< 0.05 was used. Tables were used to present the examined data.

## Results

### Phytochemical screening

The presence of alkaloids, tannins, flavonoids, and saponins were found in the ME and the solvent fractions of the leaves of *R. prinoides* (**Table 1**).

**Table 1:** Preliminary phytochemical screening of ME and solvent fractions of the leaves *R. prinoides*.

| Secondary metabolite | Methanol crude extract | n-butanol fraction | Chloroform fraction | Pet. ether fraction |
| --- | --- | --- | --- | --- |
| **Saponin** | - | + | – | – |
| **Terpinoids** | + | + | + | + |
| **Tannins** | + | + | + | - |
| **Flavonoid** | + | + | + | + |
| **Glycosides** | + | + | + | + |
| **Steroids** | + | + | – | + |
| **Alkaloids** | + | + | + | – |

(+, Present); (-, Absent)

### Acute Toxicity Test

Physical and behavioral abnormalities, as well as mortality, were not seen in the acute toxicity test with ME and the solvent fractions of the leaves of *R. prinoides* within 24 hours and for the next 14 days. The oral LD50 of the ME of *R. prinoides* and the fractions was larger than 2000 mg/kg in mice, confirming the “Limit Test” of OECD guideline 425 [9]. As a result, dosages of 100, 200, and 400 mg/kg for the ME and solvent fractions of the leaves of *R. prinoides* were determined and employed in the experiment.

### The effect of *Rhamnus prinoides* crude extract and solvent fractions on castor oil-induced diarrhea

During the four hours observation period, all animals in the control group had either wet stool or watery diarrhea. The ME of *R. prinoides*, produced a substantial decrease in the onset, the number and weight of wet and total stools at the dose of 400 mg/kg (p<0.01) which was comparable with loperamide. At the dose of 200 mg/kg ME, there was a significant decrease in the number and weight of wet and total feces (p<0.01), although the delay in the onset of diarrhea was unable to achieve a meaningful level. Moreover, the ME at the tested dose of 100 mg/kg did not show a significant reduction in all parameters used to determine antidiarrheal activity (**Table 2**). Furthermore, dose dependent percentage of inhibition of diarrhea was observed by ME of *R. prinoides:* 45.28% (R^2^=0.679), 66.41% (R^2^=0.869; p<0.01), and 79.71% (R^2^=0.953; p<0.01) for 100, 200, and 400 mg/kg respectively (**Table 2**). On the other hand, the ME of *R. prinoides* at the test doses of 200 mg/kg was unable to reach a significant level in delaying the onset of diarrhea. Percentages of inhibition for both the watery content and total weight of wet stools in comparison to negative controls and comparison among different doses were determined. When compared to the negative control, both 200 mg/kg and 400 mg/kg doses of the ME of *R. prinoides* revealed in a significant decrease in both the total weight of wet and watery content of the stool (p<0.01). When compared to the negative control, the 200 mg/kg dose, however, only showed a substantial decrease in the water content of the stools. When compared to 100 mg/kg, the dose of 400 mg/kg showed a significant decrease in both the total weight of wet and watery content of the stool (p<0.01).

**Table 2:**
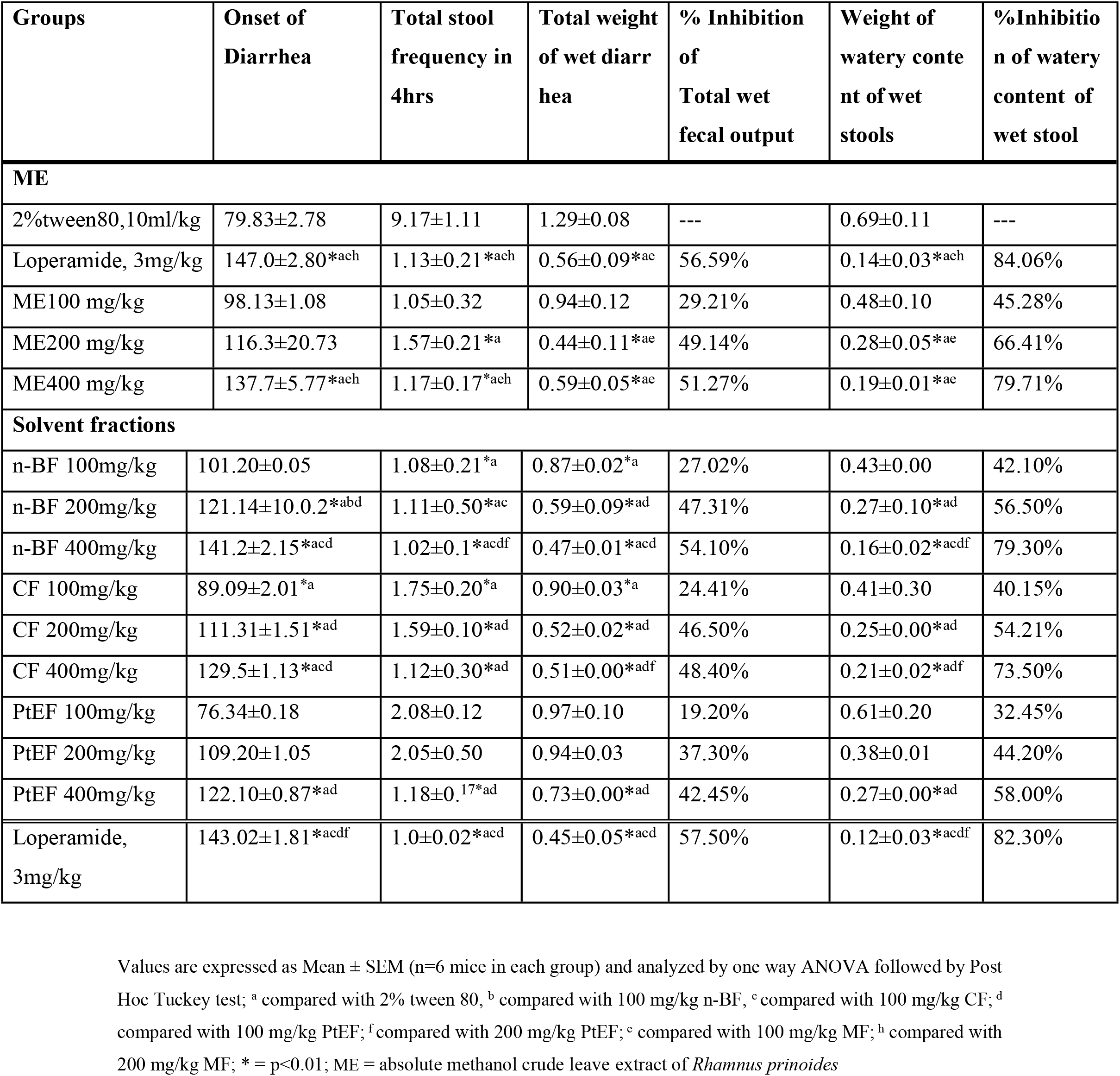
The effect of ME and solvent fractions of the leaves of *R. prinoides* on castor oil induced diarrhea model in mice.

Among the solvent fractions the leaves of *R. prinoides*, n-BF significantly delayed the onset of time of diarrheal and reduced the frequency of defecation dose dependently: 100 mg/kg (R^2^=0.801) 200 mg/kg (R^2^=0.877) and, 400 mg/kg (R^2^=0.976; p<0.01). Data obtained from the current study revealed that the percentage of inhibition of diarrhea as compared with the negative control were 56.50% and 79.30% (p<0.01) at doses of 200 and 400 mg/kg n-BF respectively (**Table 2**). Likewise, the chloroform fraction revealed dose dependent reduction of the frequency and the weight of wet and total stool at the tested dose of 100 mg/kg (R^2^=0.976), 200 mg/kg (R^2^=0.867), and, 400 mg/kg (R^2^=0.968 p<0.01) compared with the negative control at the stated doses of 200 and 400 mg/kg respectively. In contrast, the PtEF of *R. prinoides* lack a substantial decrease in the frequency, number, and weight of (the wet and total stool) at the tested doses of 100 and 200 mg/kg as compared with the negative control (**Table 2**). Additionally, the induction of defecation was only marginally delayed by the PtEF dosages of 100 and 200 mg/kg compared to the negative control. Data from the current study revealed that a significant difference in the antidiarrheal evaluation parameters was observed among different treatment groups. For instance, n-BF (at 200 and 400 mg/kg) significantly reduced the frequency, number, and weight of both wet and total stool compared with 100 and 200 mg/kg of PtEF of the leaves of *R. prinoides*. Similarly, the dose of 400 mg/kg CF showed a delay in induction of diarrhea and reduction of the frequency, number, and weight of both wet and total stool compared with 100 and 200 mg/kg PtEF but the magnitude of diarrhea inhibition is lower than the n-BF (**Table 2**). Moreover, the 100 and 200 mg/kg of PtEF lack a significant reduction in all parameters even compared with the negative control (**Table 2**). There was no significant difference between the standard and 400 mg/kg ME of *R. prinoides* and n-BF, but significant differences among doses of ME of *R. prinoides* and PtEF were examined (**Table 2**).

### The effect of ME and solvent fractions of *R. prinoides* on castor oil-induced gastrointestinal motility

As described in **Table 3** the ME of *R. prinoides* showed a significant percentage reduction in intestinal motility and the effect was dos dependent at all the tested doses (R^2^ = 0.682; 42.81%; p<0.01), (R^2^ = 0.846; 56.47%; p<0.01), and (R^2^ = 0.927; 69.23%; p<0.01), for 100, 200, and 400 mg/kg respectively) as compared to the negative control against the castor oil-induced motility. When compared to the test doses of 100 and 200 mg/kg, the dose of 400 mg/kg of the ME of *R. prinoides* showed a significant anti-motility effect (69.23%, p<0.01) which is comparable with loperamide 3 mg/kg (71.18%, p<0.01) (**Table 3**).

**Table 3:**
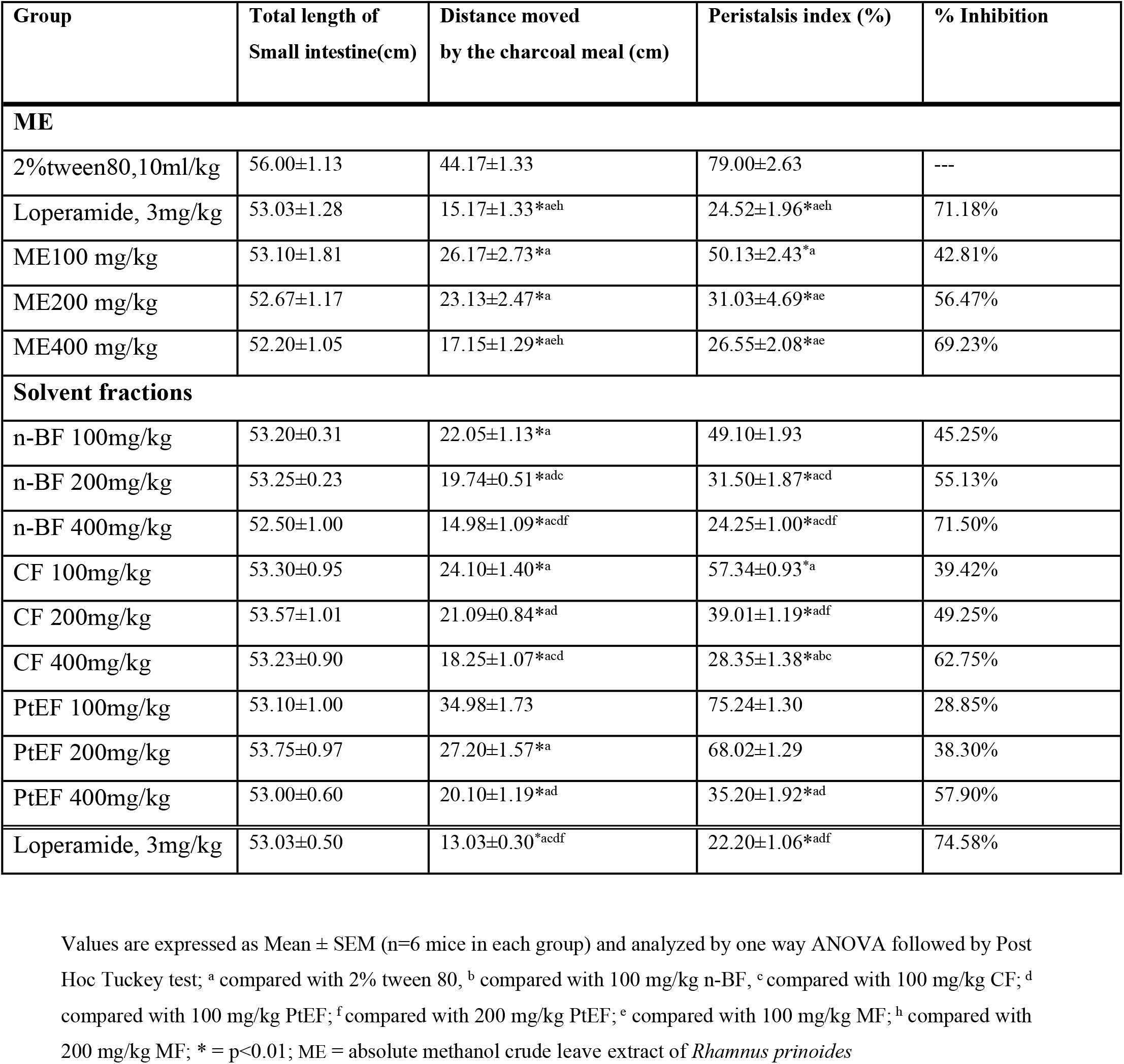
The effect of ME and solvent fractions of the leaves of *R. prinoides* on castor oil induced gastrointestinal motility in mice.

The data from the present study revealed a dose dependent reduction of the intestinal motility at all the tested doses n-BF (R^2^ = 0.781; p<0.01), (R^2^ = 0.901; p<0.01), and (R^2^ = 0.991; p<0.01) and CF ((R^2^ = 0.732; p<0.01), (R^2^ = 0.899; p<0.01), and (R^2^ = 0.956; p<0.01) correspondingly to 100, 200, and 400 mg/kg. However, the petroleum ether fraction still showed a significant reduction in intestinal motility only at the dose of 400 mg/kg as compared with the negative control (p<0.01; **Table 3**). As compared with 100 and 200 mg/kg doses of the petroleum ether fraction, the dose of 400 mg/kg n-BF revealed a significant reduction (71.50%; p<0.01) of intestinal motility which is comparable with the standard (74.58%; p<0.01).

### The effect of ME and fractions *R. prinoides* on castor oil induce enteropooling

In this antidiarrheal model, the ME of *R. prinoide* demonstrated a dose-dependent reduction in both mean weight (R^2^ = 0.676; 32.34%; p<0.01), (R^2^ = 0.798; 40.31%; p<0.01), (R^2^ = 0.899; 56.32%; p<0.01) and volume of the small intestinal content (R^2^ = 0.667; 27.07%; p<0.01), (R^2^ = 0.801; 48.02%; p<0.01), (R^2^ = 0.925; 57.05%; p<0.01) at 100, 200, and 400 mg/kg respectively (**Table 4**). Additionally, there was a significant difference in terms of volume of intestinal fluid and weight of intestinal contents among the three test doses of the ME *R. prinoide*. For instance, the dose of 400 mg/kg showed a significant reduction in both the average weight and volume of small intestine content compared with 100 and 200 mg/kg doses (p<0.01). This dose was found to have comparable antidiarrheal activity with 3 mg/kg loperamide (57.05%; p<0.01).

**Table 4:**
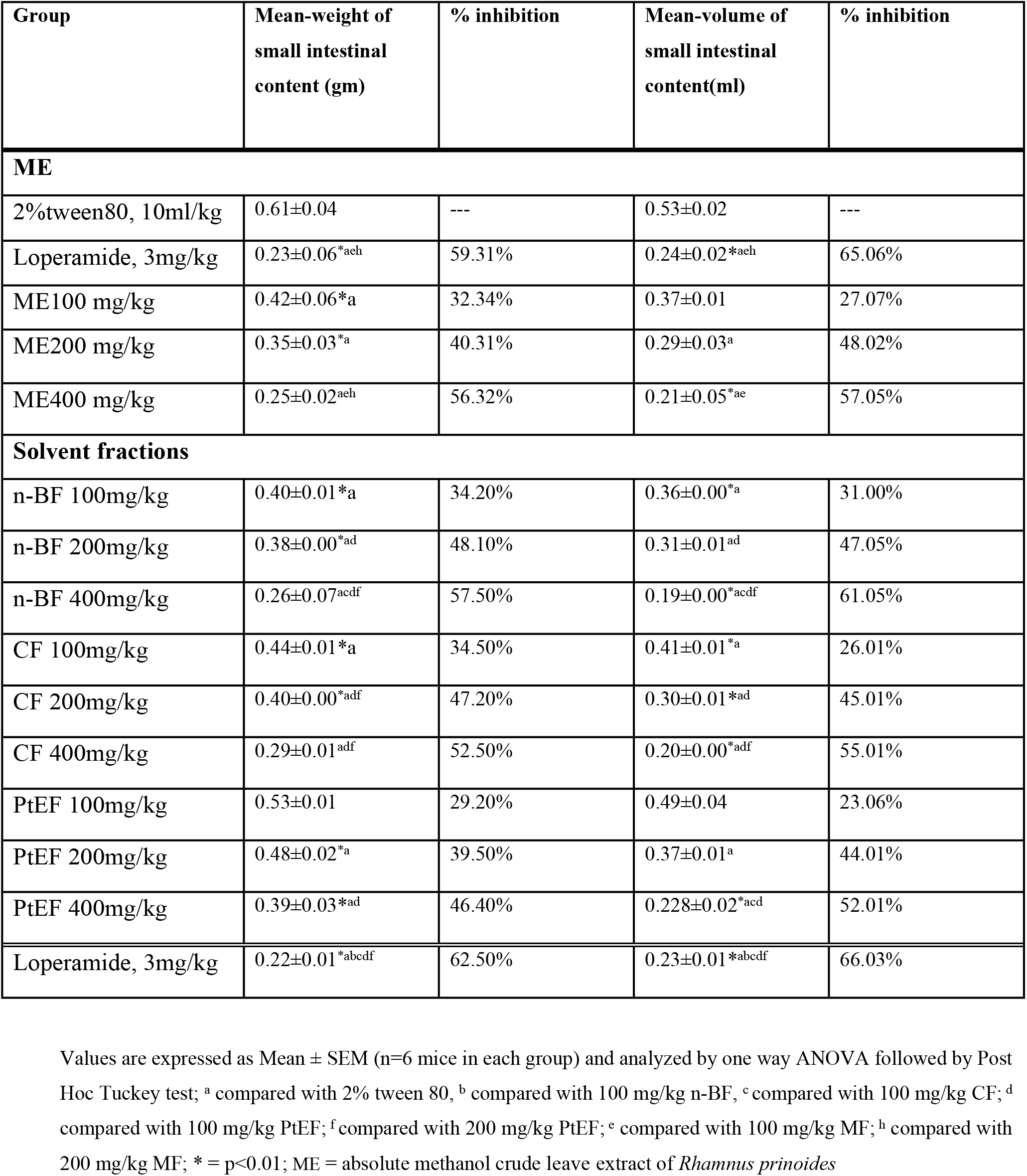
The effect of ME and solvent fractions of the leaves of *R. prinoides* on castor oil induced entropooling in mice.

The n-BF of the leaves of *R. prinoide* reduced the weight (R^2^ = 0.765; 34.20%; p<0.01), (R^2^ = 0.884; 48.10%; p<0.01), (R^2^ = 0.999; 57.50%; p<0.01) and volumes (R^2^ = 0.716; 31.00%; p<0.01), (R^2^ = 0.882; 47.05%; p<0.01), (R^2^ = 0.998; 61.05%; p<0.01) of the intestinal content at the tested of 100, 200, and 400 mg/kg respectively. As a result, it showed a comparable level of inhibition for the volume and weight of intestinal contents. In contrast, the PtEF lacks a significant reduction of the parameters used to evaluate the antidiarrheal activity by this model at the tested dose of 100 mg/kg compared with the negative control. Conversely, the percent of inhibition of the intestinal volume was significantly increased with 400 mg/kg petroleum ether fraction as compared with the negative control and 100 mg/kg dose of petroleum ether fraction (52.01%; p<0.01).

### The *in-vivo* Anti-diarrheal Index

Three parameters were used to estimate the anti-diarrheal index or protective effect of the ME and solvent fractions of *R. prinoide*. These include delay in defecation, gut meal travel distance, and purging frequency in the number of wet stools. For instance, the *in-vivo* anti-diarrheal index values were recorded high at the maximal dose (400 mg/kg) of the ME and each fraction of *Rhamnus prinoide*. ADI values are 53.24%, 64.40%, 57.50%, 53.90% for the ME, n-BF, CF, PtEF fractions respectively (**Table 5**). Moreover, the ADI values of n-BF (64.40%; 400 mg/kg) were comparable with the 3 mg/kg loperamide drug (65.90%).

**Table 5:** *In-Vivo* ADI of the ME and solvent fractions of *R. prinoides*.

| Test agents | Dose administered | Delay in defecation<br>(time of onset in<br>minute, Dfreq %) | Gut meal travel<br>distance, (Gmeq<br>%) | Reduction in<br>Intestinal fluid<br>accumulation (%) | Anti-diarrheal<br>index(ADI) |
| --- | --- | --- | --- | --- | --- |
| **ME** | ME 100mg/kg | 20.13% | 45.21% | 35.20% | 32.47% |
|  | ME 200mg/kg | 36.11% | 50.24% | 52.23% | 43.14% |
|  | ME 400mg/kg | 52.10% | 51.27% | 55.26% | 53.24% |
| **Loperamide** | 3mg/kg | 57.69% | 62.26% | 60.06% | 63.21% |
| **Solvent**<br>**fractions** | n-BF 100mg/kg | 21.20% | 46.50% | 37.30% | 35.13% |
|  | n-BF 200mg/kg | 38.50% | 52.20% | 54.40% | 48.24% |
|  | n-BF 400mg/kg | 64.30% | 59.40% | 62.60% | 64.40% |
|  | CF 100mg/kg | 20.10% | 45.50% | 34.30% | 31.40% |
|  | CF 200mg/kg | 35.20% | 49.50% | 50.10% | 45.20% |
|  | CF 400mg/kg | 56.40% | 53.30% | 52.26% | 57.50% |
|  | PtEF 100mg/kg | 19.50% | 40.50% | 30.10% | 27.30% |
|  | PtEF 200mg/kg | 30.20% | 47.40% | 39.20% | **36.50%** |
|  | PtEF 400mg/kg | 50.25% | 48.25% | 51.30% | **52.30%** |
| **Loperamide** | 3mg/kg | 67.20% | 63.50% | 62.30% | **65.90%** |

## Discussion

The purpose of this study was to determine the *in-vivo* anti-diarrheal activity of the absolute methanol leave crude extract (ME) and solvent fractions of *Rhamnus prinoide*, as well as the likely underlying mechanism. In the models employed, the plant was found to have anti-diarrheal activity.

Castor oil’s induced diarrheal has been studied using a variety of methods. These include blocking the Na+/K+-ATPase function by lowering normal fluid absorption [16] and stimulating the release of inflammatory mediators by preventing the reabsorption of NaCl and water [17]. Both the secretory and abnormal motility diarrhea are included in this model [18]. As a result, the use of castor oil as a diarrhea inducer is reasonable because it mimics the abnormal processes and allows quantifiable changes in the number of feces, intestinal transit, and enteropooling to be examined.

The standard drug, loperamide hydrochloride, not only regulates the gastrointestinal tract but also slows peristalsis across the small intestine. Nowadays, loperamide is commonly utilized in several diarrheal models to study the antidiarrheal properties of diverse experimental plants. This is due to its proven antisecretory and antimotility features [19].

In this study, the ME of *R. prinoides* showed anti-diarrheal activity by reducing castor oil-induced diarrhea in all of the models employed. The presence of various phytochemicals in the ME and solvent fractions of *R. prinoides* is the most reasonable explanation (**Table 1**). Antioxidant features were found in both flavonoids and phenolic compounds [20], which are likely to be responsible for their antidiarrheal action. These phytochemicals may work by inhibiting the enzymes that mediate arachidonic acid metabolism; hence lowering castor oil-induced fluid production [21]. In addition, phytochemical compounds such as tannins and saponins have an anti-diarrheal effect [22].

In the current investigation, there was a significant decrease (p<0.01) in the number and weight of both wet and watery fecal matter, as well as a delay in the onset of diarrhea. The effect was dose-dependently increased. This is comparable with the previous studies on several plants, in which their extracts found to have an antidiarrheal effect that was dose-dependent [23].

To ascertain the phytochemicals responsible for the antidiarrheal activity, the ME was further fractionated using solvents of different polarities via successive liquid-liquid fractionation. The maximal dose of all the fractions (n-butanol, chloroform, and petroleum ether) of *R. prinoides* significantly reduced all parameters in all diarrheal models employed with the n-BF being the most active. Furthermore, at the dose of 200 mg/kg, the n-BF, and CF fractions produced a significant reduction in both the total fecal output and the weight of wet and total defection as compared with the negative control (p<0.01). When compared to the negative control, the PtEF does not, however, significantly lower all of the antidiarrheal model’s aforementioned parameters. This may be due to quantitative and qualitative differences in the phytochemicals among the solvent fractions. For instance, the insignificant antidiarrheal effect by the petroleum ether fraction might be due to the low number and concentration of the phytoconstituents in the fraction. Another reason might be due to the differential distribution of the active phytochemicals to the n-BF and CF fractions that creates concentration variation (**Table-1**). Besides, the reduction of diarrheal parameters might be due to the polar nature of the phytochemicals in the n-BF. In general, findings from this study showed that n-BF had comparable to loperamide anti-diarrheal effects, but the effects were higher than that of the ME of *R. prinoides*. Previous reports on anti-diarrheal ethnomedicinal plants support the present study. Ethnomedicinal antidiarrheal herbs, like Eremomastax speciosa and Xylocarpus granatum, have been shown to decrease the amount of moist feces, intestinal motility, and delay the onset of diarrhea. [24, 25]. The overall order of efficacy in inhibition of diarrhea in all models is n-BF>CF>PtEF.

Because castor oil causes diarrhea by inhibiting fluid and electrolyte absorption and thus causing intestinal peristalsis, it’s a good idea to avoid it [26]. The ability of the ME and solvent fractions of *R. prinoides* to enhance fluid and electrolyte absorption through the gastrointestinal tract could be one of the causes of anti-diarrheal activity.

Additionally, following administration of the ME and solvent fractions of *R. prinoides*, there was a significantly (p<0.01) delayed onset of diarrhea and decreased fecal matter frequency (number of wet feces), suggesting their antidiarrheal effect at the test dosages used. Numerous investigations suggest that the antidiarrheal properties of the solvent and ME fractions may be attributed to phytochemicals like alkaloids, tannins, saponins, phenols, terpenoids, and flavonoids. A decrease in total fecal matter, including wet and watery content, suggests that the anti-diarrheal activity of the ME and the viable solvent fractions of the *R. prinoides* may be mediated by an antisecretory mechanism. Furthermore, the anti-diarrheal efficacy of the ME and the fractions could be attributed to its inhibitory action on nitric oxide (NO), and platelet-activating factors production and delay diarrhea caused by castor oil [27–29].

The standard drug (loperamide) used in the present study is an effective and extensively used anti-diarrheal drug which works by decreasing intestinal motility and/or inhibiting intestinal secretions [28]. Therefore, for additional validation of the potential antidiarrheal action of *R. prinoides*, the ME and solvents fraction were tested on intestinal motility and enteropooling models.

Enteropooling in mice is the technique used to evaluate the effect of investigational drugs on intestinal secretions. The study was also extended to evaluate the anti-enteropooling effect to further elucidate the mode of antidiarrheal action. When compared with the negative control, the ME (400 mg/kg), n-BF (200 and 400 mg/kg), and CF (400 mg/kg) of *R. prinoides* significantly blocked intestinal fluid collection, and the weight of intestinal content in castor oil-induced enteropooling (p<0.01). The ME and n-BF of *R. prinoides* have a comparable effect on castor oil-induced fluid buildup at their maximum dose (400 mg/kg; p<0.01). Moreover, this finding supports the idea that the *R. prinoides* ME and n-BF have an anti-enteropooling action at the dose employed for the experiment. This could be attributed to the active secondary metabolites in the ME and the n-BF which reduce and prevents fluid buildup in the gastrointestinal tract. Among the solvent fractions, the n-butanol and chloroform fractions showed a significant antimotility and antisecretory effect at all tested doses. However, the highest percentage of inhibition was recorded from the n-BF (57.50, 61.05% for mean weight and volume of intestinal contents respectively) which is comparable with the standard. Furthermore, phytochemicals such as terpenoids, steroids, flavonoids, tannins, terpenoids, flavonoids, and steroids [30–32] might inhibit the prostaglandin E2 production, which was a key factor in the activation of intestinal secretions by causing the secretion of water and electrolytes [33]. Tannins also limit fluid outflow by inhibiting CFTR and CaCC, as well as causing a protein-precipitating response in the gastrointestinal mucosa [34], making the mucosa more resistant to chemical changes [29].

The small intestine is extrinsically innervated by the autonomous nervous system [35]. For instance, neurotransmitters like acetylcholine and vasoactive intestinal peptides from the parasympathetic system activate intestinal homeostasis, whereas the sympathetic system uses 2 adrenergic substances like enkephalins and somatostatins to increase intestinal absorption. Phytochemicals from plants, such as flavonoids, may activate 2 adrenoreceptors in the gastrointestinal tract’s absorptive cells [36]. Transport of fluid over the epithelium of the gastrointestinal tract is also controlled by managing aquaporin (AQP) type water channels, in addition to electrolyte movement. Tannins were discovered to decrease the production of particular AQPs by downregulating different kinases. AQP inhibits the protein kinase signaling pathway, which accounts for some of the anti-secretory and thus anti-diarrheal effects [37]. As a result, the anti-secretory effect of the ME and solvent fractions of *R. prinoides* is most likely related to the presence of phytochemicals and their synergistic effects. With this in mind, the ME, n-BF, and CF are more likely to reduce diarrhea in this study by either boosting fluid or electrolyte reabsorption through sympathetic activation or inhibiting fluid release into the colon through altered parasympathetic activity. However, n-BF showed a superior antisecretory effect as compared with ME. This may be associated with the differential distribution of the phytochemicals in the n-BF [31]. On the contrary, antisecretory action was reduced in the PtEF as compared with ME, n-BF, and CF of *R. prinoides* might be due to the absence of phytochemicals responsible for antisecretory action (**Table 1**).

One strategy to increase the production of diarrhea is to increase intestinal motility. The study used charcoal meal as a marker to test the antimotility activity of the ME of *R. prinoides*. The sympathetic and parasympathetic nerves are the main regulators of gastrointestinal tract motility, with the latter being the most important. Parasympathetic nerve activation increases the intestinal transit, while sympathetic nerve stimulation inhibits intestinal transit [35]. Because of its anticholinergic, antihistamine, and prostaglandin blocking actions, loperamide, a common medicine, was known to slow the movement of the charcoal meal [38]. In addition to cholinergic receptors, stimulation of 2 adrenoreceptors in the gastrointestinal tract inhibits peristalsis, reduces gastrointestinal smooth muscle contraction, improves gastric emptying, and promotes stomach mucosa protection [39, 40]. The reduction in the traveled distance could be used to explain the relaxation of intestinal smooth muscles. According to previous studies, all smooth muscle contractions are completely dependent on the presence of Ca2+, which stimulates the contractile elements and causes their relaxation, a mechanism that has been linked to the antidiarrheal activity of several medications [34]. As a result, the ME of *R. prinoides* could have caused the charcoal to cover less distance by boosting intracellular Ca2+ release.

The castor oil-induced gastrointestinal motility test revealed that the ME *Rhamnus prinoide* at the tested dose of 400 mg/kg significantly reduced the intestinal propulsive movement as compared with the negative control. When compared to normal intestinal transit, the inhibitory effect of the ME of *R. prinoides* in castor oil-induced intestinal transit was greater. The *R. prinoides* crude extract’s antimotility effect may be mediated through various mechanisms. Anti-diarrhea drugs are known for stimulating intestinal relaxation, which slows the emptying time [41], allowing for better fluid absorption [42]. The anti-diarrheal effect of the ME of *R. prinoides* can therefore be explained by the extracts’ ability to inhibit intestinal movement (**Table 3**). In other words, the crude extracts’ inhibitory effect would be higher as intestinal motility increased. The significance of this discovery shouldn’t be disregarded, as it means that the risk of constipation, which is a common side effect of most conventional medications, including loperamide, will be reduced. The presence of phytochemicals that are responsible for the anti-motility action of ME of *R. prinoides* could be linked to the presence of phytochemicals that are responsible for the anti-motility action. For instance, flavonoids isolated from *Catha edulis* have been shown to inhibit intestinal smooth muscle contraction by blocking the Ca2+ channels [43, 44]. Moreover, the active constituents like ternatin and quercetin reduced the gastrointestinal motility induced by castor oil in mice [45, 46]. Furthermore, flavonoids isolated from *Baccharis teindalensis*, such as apigenin, sakuranetin, and kaempferol also decreased the hypercontractility of gastrointestinal smooth muscles [47]. Tannins such as procyanidin-B2, epicatechin, and epigallocatechin gallate isolated from rhubarb revealed an antimotility effect by inhibiting aquaporins 2 and 3 expression [48]. According to other study, tannins isolated from the bark of *ceriops decandra* had nitric oxide scavenging and antioxidant action and exhibited a promising *in-vivo* antimotility activity [49].

The antimotility effects of the n-BF and CF fractions were both statistically significant, with the n-BF being the most active at the tested dose of 400 mg/kg (71.50%, p<0.01). This may be as a result of varied distribution of the more active phytoconstituents in the polar and less polar solvents [50]. Because of the abundant phytochemical constituents in the solvent fraction, n-BF revealed antimotility effect in a dose dependent way. Moreover, flavonoids; Luteolin and apigenin isolated from ethyl acetate fraction of *D. kotschyi* showed a significant inhibitor effect on the gastrointestinal motility activity [51]. Alkaloid fraction isolated from. *W. tinctoria* possess antimotility, and antiperistaltic activity [52]. Although the PtEF significantly reduced the distance covered by the charcoal meal, it had a little inhibitory effect on the intestinal motility, especially at the tested dose of 100 mg/kg. Additionally, in all models the maximum and minimum antidiarrheal effects of n-BF and PtEF, respectively, are consistent with previous studies [53]. This demonstrates the variation in the kind and concentration of the bioactive phytoconstituents in the fractions, the most active phytochemicals being localized in the n-BF and CF.

The ADI is a method of calculating the cumulative effects of antidiarrheal parameters such as decreased GI motility, the beginning of diarrheal stools, and fluid accumulation [46]. According to the literature, the extract’s or the fraction’s effectiveness at treating diarrhea increases with increasing ADI values [34]. In the entire diarrheal model employed, the antidiarrheal activity of the crude extract and solvent fractions of *R. prinoides* leaves was increased dose dependently: ME (R^2^=0.921), n-BF (R^2^=0.987), CF (R^2^=0.951), PtEF (R^2^=0.821). However the ADI value for n-BF at the tested dose of 400 mg/kg was higher than all tested dose of the ME of *R. prinoides* leaves. This may be the loss of the additive and synergistic effect of amongst the phytochemicals in the ME, while the abundance of concentrated bioactive phytochemical in the n-BF. Moreover, the ADI value also confirmed that the antidiarrheal activity of the n-BF of *R. prinoides* leaves was comparable to that of the standard medication. On the other hand, the ADI of PtEF of the leave of *R. prinoides* was low, suggesting minimal antidiarrheal activity in all models employed.

## Conclusion

The current findings demonstrated the anti-diarrheal properties of the ME of *R. prinoides* leaves in all the models used. Additionally, all three solvent fractions have variable magnitude of antidiarrheal activity, with n-BF being the most active fraction in the reduction of all the parameters in all aforementioned models. The inclusion of bioactive substances such terpenoids, flavonoids, tannins, alkaloids, and glycosides that act singly or together may be responsible for the antidiarrheal properties of the ME as well as fractions. Furthermore, the findings of this study indicate that phytochemicals with semi-polar to non-polar properties are more likely to be in charge of the antidiarrheal activity that was seen.

## Declarations

### Ethics approval

Ethical clearance and permission was obtained from Debre Tabor University Research and Ethical Review Committee and the approval was obtained by protocol number CHS/135/2022.

### Data Availability

All the data sets generated and analyzed during the study are included in the text.

## Acknowledgments

I would also like to thank Ethiopian Public Health Institute providing us laboratory animals; Department of Pharmacology, School of Pharmacy, University of Gondar and Debre Tabor University for allowing us to use their laboratory facilities.

## Authors’ contribution

The authors are in charge of the paper’s content and writing. TM developed the draft proposal under the supervision of. EC, AB, ZT, MM, YS, GT, GA, MA, SB, NT, TY and AE critically contributed to the conceptualization, data collection, analysis, and writing of the paper. TM, EC. AB, ZT, MM, YS, GT, GA, MA, SB, NT, TY and AE also participated in manuscript preparation, and they read and approved the final manuscript.

## References

1. Karam S, Bhavna V. An integrated approach towards Diarrhoea. Research and Reviews: Journal of AYUSH. 2012;1(2):11–33.

2. Rout SP, Choudary KA, Kar DM, Das LO, Jain A. Plants in traditional medicinal systemfuture source of new drugs. Int J Pharm Pharm Sci. 2009 Jul;1(1):1–23.

3. Prasad DR, Izam A, Khan MM. Jatropha curcas: Plant of medical benefits. Journal of medicinal plants research. 2012 Apr 16;6(14):2691–9.

4. Ashu Agbor M, Naidoo S. Ethnomedicinal plants used by traditional healers to treat oral health problems in Cameroon. Evidence-Based Complementary and Alternative Medicine. 2015 Oct 1;2015.

5. Camilleri M, Sellin JH, Barrett KE. Pathophysiology, evaluation, and management of chronic watery diarrhea. Gastroenterology. 2017 Feb 1;152 (3):515–32.

6. Alemu A, Tamiru W, Nedi T, Shibeshi W. Analgesic and anti-inflammatory effects of 80% methanol extract of Leonotis ocymifolia (burm. F.) iwarsson leaves in rodent models. Evidence-Based Complementary and Alternative Medicine. 2018 Jan 1;2018.

7. Yeshitila A. Phytochemical investigation on the leaves of leonotis ocymifolia, AAU, 2006.

8. Ahmadi-Noorbakhsh S, Mirabzadeh Ardakani E, Sadighi J, Aldavood SJ, Farajli Abbasi M, et al. Guideline for the Care and Use of Laboratory Animals in Iran. Lab animal. 2021 Nov;50(11):303–5.

9. Buschmann J. The OECD guidelines for the testing of chemicals and pesticides. Teratogenicity testing. 2013:37–56.

10. Sasidharan S, Chen Y, Saravanan D, Sundram KM, Latha LY. Extraction, isolation and characterization of bioactive compounds from plants’ extracts. African journal of traditional, complementary and alternative medicines. 2011;8(1).

11. Ayoola GA, Coker HA, Adesegun SA, Adepoju-Bello AA, Obaweya K, Ezennia EC, et al. Phytochemical screening and antioxidant activities of some selected medicinal plants used for malaria therapy in Southwestern Nigeria. Tropical Journal of Pharmaceutical Research. 2008 Sep 11;7(3):1019–24.

12. Umer S, Tekewe A, Kebede N. Antidiarrhoeal and antimicrobial activity of Calpurnia aurea leaf extract. BMC complementary and alternative medicine. 2013 Dec;13(1):1–5

13. Oghenesuvwe EE, Tedwins EJ, Obiora IS, Lotanna AD, Treasure UN, Ugochukwu OM, Emmanuel IE. Preclinical screening techniques for anti-diarrheal drugs: a comprehensive review. American Journal of Physiology. 2018;7(2):61–74.

14. Amuzat A, Ndatsu Y, Adisa M, Sulaiman R, Mohammed H, Yusuf A, et al. Anti-diarrhoeal Effects of Aqueous Extract of Vitex doniana Stem Bark in Castor Oil-induced Wistar Rats. Tanzania Journal of Science. 2020 Oct 30;46(3):723–32.

15. Uzuegbu UE, Mordi JC, Ovuakporaye SI, Ewhre LO. Effects of aqueous and ethanolic extracts of Tridax procumbens leaves on gastrointestinal motility and castor oil-induced diarrhoea in wistar rats. Biokemistri. 2021 Jun 10;27(1).

16. Yakubu MT, Amoniyan OD, Mohammed MO, Assin CI, Abubakar JO, Salimon SS, et al. Anti-diarrhoeal activity of aqueous extract of Cochlospermum planchonii (Hook Fx. Planch) leaves in female Wistar rats. Journal of Medicinal Plants for Economic Development. 2020 Jan 1;4(1):1–8.

17. Imam MZ, Sultana S, Akter S. Antinociceptive, antidiarrheal, and neuropharmacological activities of Barringtonia acutangula. Pharmaceutical Biology. 2012 Sep 1;50(9):1078–84.

18. Achenef B. a Thesis Paper Submitted To the Department of Pharmacology and Presented in Partial Fulfillment of the Requirements for the Master’s Degree of. 2019;

19. Salgado H, Roncari AF, Moreira RR. Antidiarrhoeal effects of Mikania glomerata Spreng.(Asteraceae) leaf extract in mice. Revista Brasileira de Farmacognosia. 2005 Sep;15(3):205–8.

20. El-Mostafa K, El Kharrassi Y, Badreddine A, Andreoletti P, Vamecq J, El Kebbaj MH, Latruffe N, Lizard G, Nasser B, Cherkaoui-Malki M. Nopal cactus (Opuntia ficus-indica) as a source of bioactive compounds for nutrition, health and disease. Molecules. 2014 Sep 17;19(9):14879–901.

21. Jalilzadeh-Amin G, Maham M. The application of 1, 8-cineole, a terpenoid oxide present in medicinal plants, inhibits castor oil-induced diarrhea in rats. Pharmaceutical Biology. 2015 Apr 3;53(4):594–9.

22. Muthukumaran P, Saraswathy N, Aswitha V, Balan R, Gokhul VB, Indumathi P, et al. Assessment of total phenolic, flavonoid, tannin content and phytochemical screening of leaf and flower extracts from Peltophorum pterocarpum (DC.) Backer ex K. Heyne: a comparative study. Pharmacognosy Journal. 2016;8(2).

23. Krueger RJ. Drugs of Natural Origin. A Textbook of Pharmacognosy. By Gunnar Samuelson. Swedish Pharmaceutical Press, Stockholm. 620 pp. 17×25 cm. $70.00. ISBN 91-9743-184-2.

24. Oben JE, Sheila EA, Gabrie AA, Musoro DF. Effect of eremomastax speciosa on experimantal diarrhoea. African Journal of Traditional, Complementary and Alternative Medicines. 2006 Jan 12;3(1):95–100.

25. Rouf R, Uddin SJ, Shilpi JA, Alamgir M. Assessment of antidiarrhoeal activity of the methanol extract of Xylocarpus granatum bark in mice model. Journal of Ethnopharmacology. 2007 Feb 12;109(3):539–42.

26. Nu R, Aslam K, Urooj F, Mahrukh A, Nawal AM, Massarani SA, et al. Presence of laxative and antidiarrheal activities in Periploca aphylla: A Saudi medicinal plant. International Journal of Pharmacology. 2013;9(3):190.

27. Tangpu V, Yadav AK. Antidiarrhoeal activity of Rhus javanica ripen fruit extract in albino mice. Fitoterapia. 2004 Jan 1;75(1):39–44.

28. Adzu B, Amos S, Amizan MB, Gamaniel K. Evaluation of the antidiarrhoeal effects of Zizyphus spina-christi stem bark in rats. Acta tropica. 2003 Jul 1;87(2):245–50.

29. Balaji G, Chalamaiah M, Ramesh B, Reddy YA. Antidiarrhoeal activity of ethanol and aqueous extracts of Carum copticum seeds in experimental rats. Asian Pacific Journal of Tropical Biomedicine. 2012 Feb 1;2(2):S1151–5.

30. Malik AR, Wani AH, Bhat MY, Parveen S. Ethnomycological knowledge of some wild mushrooms of northern districts of Jammu and Kashmir, India. Asian Journal of Pharmaceutical and Clinical Research. 2017 Sep 1:399–405.

31. Hämäläinen M, Nieminen R, Asmawi MZ, Vuorela P, Vapaatalo H, Moilanen E. Effects of flavonoids on prostaglandin E2 production and on COX-2 and mPGES-1 expressions in activated macrophages. Planta Medica. 2011 Sep;77(13):1504–11.

32. Awad AB, Toczek J, Fink CS. Phytosterols decrease prostaglandin release in cultured P388D1/MAB macrophages. Prostaglandins, Leukotrienes and Essential Fatty Acids. 2004 Jun 1;70(6):511–20.

33. Pierce NF, Carpenter Jr CC, Elliott HL, Greenough III WB. Effects of prostaglandins, theophylline, and cholera exotoxin upon transmucosal water and electrolyte movement in the canine jejunum. Gastroenterology. 1971 Jan 1;60(1):22–32.

34. Araj-Khodaei M, Noorbala A.A, Yarani R, Emadi F, Emaratkar E, Faghihzadeh S, et al. A double-blind, randomized pilot study for comparison of Melissa officinalis L. and Lavandula angustifolia Mill. with Fluoxetine for the treatment of depression. BMC complementary medicine and therapies, 2020; 20(1), pp.1–9.

35. Tortora G. J, & Derrickson B. Principles of Anatomy and Physiology. The digestive system; Neural innervation of the gastrointestinal system. 2009; 925–926

36. Araj-Khodaei M, Noorbala AA, Yarani R, Emadi F, Emaratkar E, Faghihzadeh S, et al. A double-blind, randomized pilot study for comparison of Melissa officinalis L. and Lavandula angustifolia Mill. with Fluoxetine for the treatment of depression. BMC complementary medicine and therapies. 2020 Dec;20(1):1–9.

37. Liu C, Zheng Y, Xu W, Wang H, Lin N. Rhubarb tannins extract inhibits the expression of aquaporins 2 and 3 in magnesium sulphate-induced diarrhoea model. BioMed research international. 2014 Oct;2014.

38. Karim SM, Adaikan PG. The effect of loperamide on prostaglandin induced diarrhoea in rat and man. Prostaglandins. 1977 Feb 1;13(2):321–31.

39. Beserra FP, de Cássia Santos R, Périco LL, Rodrigues VP, de Almeida Kiguti LR, Saldanha LL, et al. Cissus sicyoides: pharmacological mechanisms involved in the antiinflammatory and antidiarrheal activities. International journal of molecular sciences. 2016 Feb;17(2):149.

40. Suleyman H. The role of alpha-2 adrenergic receptors in anti-ulcer activity. The Eurasian Journal of Medicine. 2012 Apr;44(1):43. Crombie H, Gallagher R, Hall V. Assessment and management of diarrhoea. Nursing Times. 2013 Jul 1;109(30):22–4.WGO. (2013). Acute diarrhea in adults and children. Clinical Gatereology guideline, 47. Panda SK, Patra N, Sahoo G, Bastia AK, Dutta SK. Anti-diarrheal activities of medicinal plants of Similipal Biosphere Reserve, Odisha, India. International Journal of Medicinal and Aromatic Plants. 2012;2(1):123–34. Arslan H, Inci EK, Azap OK, Karakayali H, Torgay A, Haberal M. Etiologic agents of diarrhea in solid organ recipients. Transplant Infectious Disease. 2007 Dec;9 (4):270–5. Arifeen S, Black RE, Antelman G, Baqui A, Caulfield L, Becker S. Exclusive breastfeeding reduces acute respiratory infection and diarrhea deaths among infants in Dhaka slums. Pediatrics. 2001 Oct 1;108 (4):e67-. Yacob T, Shibeshi W, Nedi T. Antidiarrheal activity of 80% methanol extract of the aerial part of Ajuga remota Benth (Lamiaceae) in mice. BMC complementary and alternative medicine. 2016 Dec;16(1):1–8

41. Vidari G, Vita Finzi P, Zaruelo A. Antiulcer andantidiarrhoeic effect of Baccharis teindalensis. Pharm Biol 41: 2003; 405–411

42. Chunfang L, Yanfang Z, Wen X, Hui W, Na L. Rhubarb Tannins Extract Inhibits the Expression of Aquaporins 2 and 3 in Magnesium Sulphate-Induced Diarrhoea Model. Hindawi Publishing Corporation BioMed Research International Volume 2014, Article ID 619465, 14 pages

43. Hemayet H, Musfizur H, Ismet-Ara J, Ishrat N, Amirul I. Antidiarrhoeal Activity, Nitric Oxide Scavenging and Total Tannin Content From The Bark Of Ceriops Decandra (Griff.) Ding Hou. Hossain et al. IJPSR 2012; Vol. 3(5): 1306–1311

44. Misganaw D, Engidawork E, and Nedi T. Evaluation of the anti-malarial activity of crude extract and solvent fractions of the leaves of Olea europaea (Oleaceae) in mice. BMC Complementary and Alternative Medicine (2019) 19:171

45. Sadraei H, Ghanadian SM, Moazeni S. Inhibitory effect of hydroalcoholic and flavonoids extracts of Dracocephalum kotschyi, and its components luteolin, apigenin and apigenin-4’-galactoside on intestinal transit in mice. J Herbmed Pharmacol. 2018;9(1):8–13.

46. Papiya B, Avtar CR. Antidiarrheal and Antispasmodic Activity Of Wrightia Tinctoria Bark And Its Steroidal Alkaloid Fraction. Pharmacologyonline 2009; 3: 298–310

47. Molla M, Gemeda N, Abay SM. Investigating potential modes of actions of Mimusops kummel fruit extract and solvent fractions for their antidiarrheal activities in mice. Evidence-Based Complementary and Alternative Medicine. 2017 May 9;2017.

